# A tricomponent mRNA pertussis vaccine elicits superior protective immunity over licensed acellular pertussis vaccine against emerging antibiotic-resistant variants

**DOI:** 10.64898/2026.11.26.769370

**Authors:** Wei Zhang, Jie Che, Guan Wang, Jingyi Wang, Si Qin, Hanying Dai, Junchi Ma, Junyue Meng, Xianyu Shao, Kun Guo, Zhenhua Di, Qiongyu An, Pinting Xiao, Shentian Zhuang, Xinghua Gao, Mingyun Shen, Yong Yang, Xiao Ma, Liyong Yuan, Maojun Zhang, Lichan Wang, Zhujun Shao, Ang Lin

## Abstract

Pertussis remains a major global health burden and has been resurging rapidly over the past decade. The causes of pertussis re-emergence are multifactorial, including vaccine-induced pathogen evolution, antimicrobial resistance, and crucially suboptimal protective immunity elicited by the widely used acellular pertussis vaccine (aPV). In this study, we developed a tricomponent mRNA pertussis vaccine formulated with lipid nanoparticle (LNP), designated mPervax, which encodes inactive pertussis toxin subunit 1 (PTS1), truncated filamentous hemagglutinin (FHA) and pertactin (PRN). Compared to a commercialized Tetanus-diptheria-pertussis vaccine Boostrix, mPervax induced stronger and broader innate immune activation, superior antigen-specific Th1-type CD4^+^ T and Tfh cell responses, and robust *B. pertussis*-specific antibody responses with remarkably increased magnitude and functionality. In addition, mPervax conferred an efficient maternal antibody-mediated protection in offspring mice and reshaped the immune profile established by primary Boostrix immunization towards a Th1-phenotype when used as a heterologous booster. Furthermore, upon bacterial challenge, mPervax outperformed Boostrix not only against the WHO reference strain 18323, but strikingly provided effective protection against emerging antibiotic-resistant clinical isolates that are deficient in PRN and/or FHA. Collectively, these data demonstrate that the mRNA pertussis vaccine is superior to Boostrix in multiple contexts, and therefore holds promise for next-generation pertussis vaccine development.

## Introduction

Pertussis is an acute, highly transmissible respiratory disease caused by *Bordetella pertussis* (*B. pertussis*), which has re-emerged in recent years despite high global vaccination coverage. A major global pertussis resurgence occurred in 2024, with nearly one million reported cases that substantially exceeded pre-pandemic levels^1–3^. The resurgence of pertussis is multifactorial and has been attributed to multiple factors, including bacterial evolution, antimicrobial resistance, improved diagnostics, and critically the suboptimal vaccine-induced immunity^4^.

Currently, two types of pertussis vaccines, whole-cell pertussis vaccine (wPV) and acellular pertussis vaccine (aPV), are used in a combined form with other components against diphtheria, tetanus and/or polio^5^. Substantial evidence has indicated that efficient protection against pertussis requires both specific antibodies and CD4^+^ T cells, and the distinctly shaped vaccine immunity determines the quality of protection^6,7^. wPV and aPV differ substantially in magnitude and functionality of the elicited vaccine responses, which has been attributed as a fundamental reason underlying pertussis resurgence^8–10^. Previous studies have indicated that wPV predominantly induces Th1 and/or Th17 T cell responses, resembling those induced by natural *B. pertussis* infection^11^. In addition, wPV-induced antibodies exhibit a more Th1-skewed phenotype that enhances bactericidal activity through Fc-mediated engagements with innate effector cells^12^. By contrast, aPV-induced vaccine responses are largely restricted to a Th2 phenotype that is less efficient at controlling *B. pertussis* infection^13^. Notably, wPV is significantly more effective than aPV in preventing nasopharyngeal colonization following *B. pertussis* infection, which is critical for blocking transmission ^11^. As a consequence, disease burden of pertussis is increasing not only in infants but also among school-aged children and adolescents, in whom infection is often asymptomatic or mild but still contributes to transmission to infants ^14^. Beyond the suboptimal immunity induced by aPV, the emergence of *B. pertussis* variants poses additional challenges to pertussis control. Under vaccine pressure, mutated variants deficient in key antigens included in vaccine formulations have emerged in multiple regions ^15,16^. Pertactin (PRN)-deficient, filamentous hemagglutinin (FHA)-deficient, and PRN/FHA double-deficient isolates have all been identified clinically^17^. In addition, macrolide-resistant *B. pertussis* (MRBP) variants have been detected worldwide and pose a significant challenge to the effectiveness of antibiotic treatment^18,19^. These challenges thus highlight the need for new or improved pertussis vaccines capable of providing broader, more efficient, and durable protection against evolved variants.

The well-established association between Th1-type anti-pertussis immunity and effective protection underscores the rationale for developing next-generation pertussis vaccine using the emerging mRNA platform. Unlike conventional protein-based subunit vaccines, we have previously shown in multiple contexts that mRNA vaccines have superior capacity to elicit comprehensive vaccine responses, including robust innate immune activation, strong Th1-polarized T cell and cytotoxic T lymphocyte (CTL) responses, as well as durable antibody responses mediating efficient bactericidal function^20–23^. In this regard, transforming protein-based aPV into mRNA counterparts may shift its immune attributes towards a phenotype favoring the control of *B. pertussis* infection and transmission. In the present study, we developed a tricomponent mRNA pertussis vaccine candidate designated mPervax, and provided an in-depth evaluation of mPervax benchmarked against the commercialized Boostrix vaccine. The data showed that mPervax is superior to Boostrix across a wide spectrum of parameters associated with vaccine efficacy, including stronger and broader innate immune activation, superior antigen-specific Th1-type CD4^+^ T and Tfh cell responses, and robust *B. pertussis*-specific antibody responses with remarkably high magnitude and functionality. In addition, mPervax conferred efficient maternal antibody-mediated protection in offspring mice and reshaped the immune profile established by primary Boostrix immunization towards a Th1-phenotype when used as a heterologous booster. Moreover, mPervax outperformed Boostrix in providing effective protection against emerging macrolide-resistant clinical variants deficient in PRN and/or FHA.

## Results

### Rational design and construction of mRNA pertussis vaccine candidates

The authorized aPVs typically contain three to five *B. pertussis* antigens including PT, FHA, PRN, fimbriae 2 and 3 (FIM2/3), which are major immunogens able to mount specific protective immunity^24^. In our study, we selected three principal antigens (PT, FHA and PRN) in their rationally modified forms as the antigen basis for mRNA vaccine development (Fig. 1a). The PT component was modified as a N-terminally truncated form of PT subunit 1 (PTS1) with two amino acid substitutions (R9K and E129G), which was intended to eliminate enzymatic toxicity while preserving immunogenicity^25^. A truncated form of FHA containing 1073–2440 residues covering the type I immunodominant domain (residues 1655–2110) was used, which was deprived of the weakly immunogenic N-terminal region and central repeat sequences that may divert vaccine immunity^26–28^. PRN was retained in its full-length form as the third antigen component in the vaccine formulation. All mRNA constructs were fused at the N-terminus to a mammalian signal peptide derived from the IgG kappa chain to direct the encoded proteins into the secretory pathway for promoted secretion (Fig. 1a). Further, using our well-established platform^20,29–32^, we encapsulated mRNA payload into lipid nanoparticle (LNP) using a classical formulation containing ionizable lipid SM102, cholesterol, DSPC, and PEG2000-DMG at molar ratios of 50: 38.5: 10: 1.5%, and generated three mono-component mRNA vaccines (Fig. 1b). Dynamic light scattering analysis showed that the three vaccine formulations had mean particle diameters of 92.07, 105.09, and 95.20 nm, respectively, and showed a polydispersity index (PDI) below 0.2, a zeta potential value between - 10 and + 0 mV, and a high encapsulation efficiency exceeding 95% (Fig. 1c).

**Figure 1.**
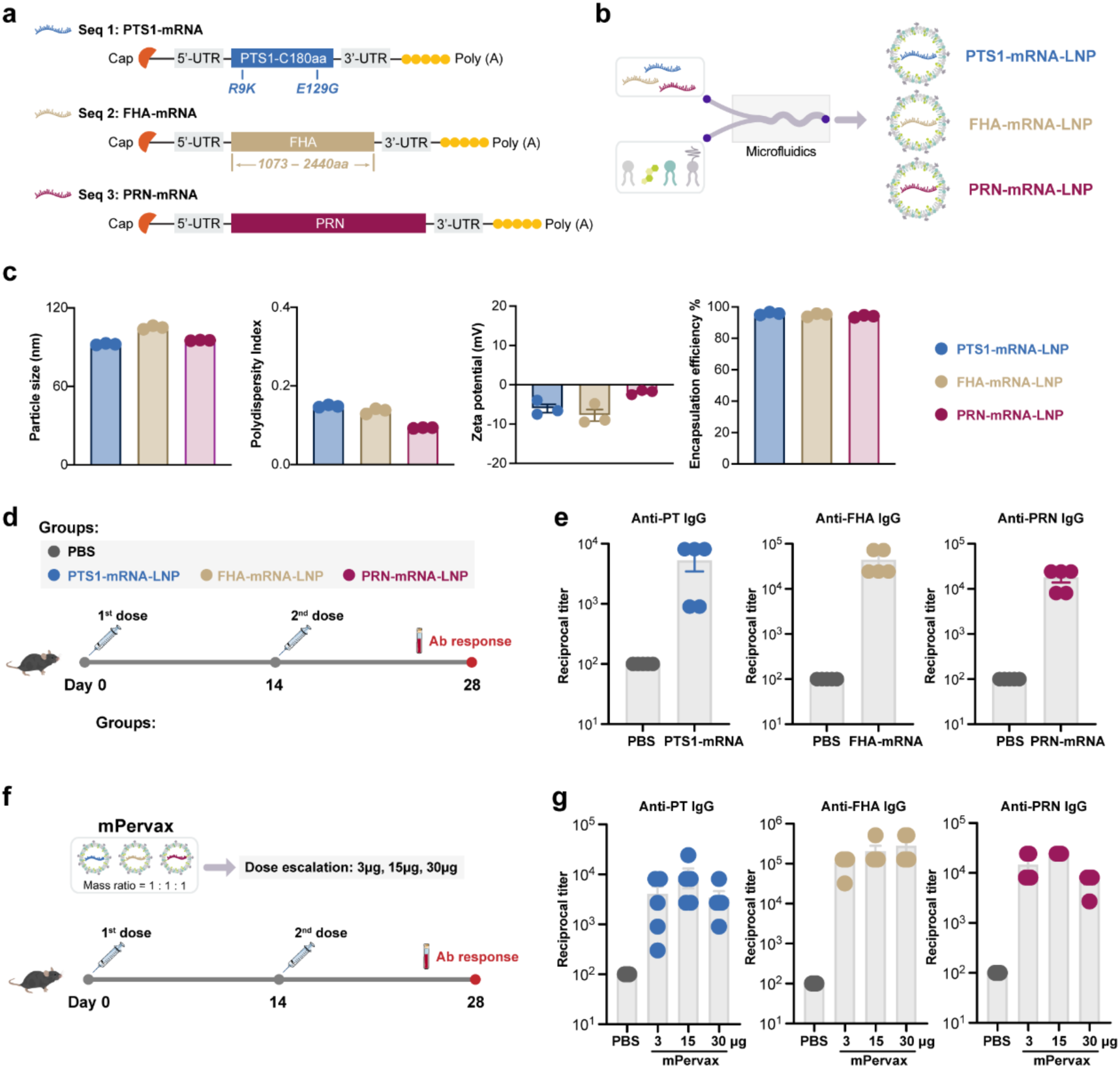
Rational design of mRNA pertussis vaccine and preliminary immunogenicity evaluation. **a-b,** Schematic design of mRNAs encoding three *B. pertussis* antigens and preparation of LNP-mRNA vaccine formulations. **c,** Characterization of LNP-mRNA formulations and major parameters are shown. **d,** Experimental design. C57BL/6 mice (n=5/group) were immunized with two doses of the three indicated mono-component vaccines at a dose of 10 μg. **e,** Antigen-specific IgG at day 14 after boost immunization was evaluated. Endpoint titers are shown. **f,** Experimental design. C57BL/6 mice (n=5/group) were immunized with escalating doses (3 μg, 15 μg, 30 μg) of mPervax vaccine at an interval of 2 weeks. **g,** Antigen-specific IgG at day 14 after boost immunization was evaluated. Endpoint titers are shown. Data are shown as mean ± SEM.

We first evaluated immunogenicity of the three mono-component mRNA vaccines by immunizing C57BL/6 mice twice at a 2-week interval (Fig. 1d). Robust levels of antigen-specific IgG against PT, FHA and PRN were detected in all immunized animals, which suggested high vaccine immunogenicity (Fig. 1e). Further, the three mono vaccine formulations were combined at a 1:1:1 mass ratio to generate a tricomponent mRNA pertussis vaccine designated as mPervax (Fig. 1f). Following a two-dose vaccination regimen, mPervax induced robust dose-escalating IgG responses across all three *B. pertussis* antigens (Fig. 1g). The response was more prominent in mice receiving 15-μg vaccination and no inter-antigen interference was observed. In addition, given the central role of anti-PT immunity in controlling pertussis^33,34^, we compared the immunogenicity of mPervax with a modified formulation containing PTS1-mRNA, FHA-mRNA and PRN-mRNA at a 2: 1: 1 mass ratio. At an equivalent 15 μg dose, the two vaccine formulations elicited a comparable level of antigen-specific IgG (Extended Data Fig. 1a-b). While interestingly, increasing the proportion of PTS1-mRNA component showed a negative effect on the generation of antigen-specific IFN-γ-producing T cells (Extended Data Fig. 1c-d).

### Innate immune profiling of mPervax in comparison with Boostrix vaccine

Vaccine-induced early innate immune activation fundamentally determines the establishment of adaptive responses^35^, we next performed an in-depth profiling of innate immune responses induced by mPervax (15 μg dose), which was benchmarked against that by a commercialized vaccine Boostrix (1/3 human dose, GSK) containing aPV component. 12 hours post injection, mPervax induced remarkably stronger activation of multiple innate cell subsets in draining lymph node (LN), including monocytes, neutrophils, conventional dendritic cells (cDCs), and plasmacytoid dendritic cells (pDCs) (Fig. 2a-b). To gain a deeper insight into the innate immune signature, we further performed RNA sequencing (RNA-seq) and transcriptomic analysis on the draining LNs. Principal component analysis (PCA) revealed markedly distinct transcriptional profiles among the three groups, with the mPervax-treated samples clustering well apart from those of the Boostrix and PBS groups (Extended Data Fig. 2a). Compared with the PBS group, Boostrix induced 86 differentially expressed genes (DEGs; 76 upregulated and 10 downregulated), which were mainly enriched for lipid and amino acid metabolism (*Ucp3, Me1, Thrsp*), whereas mPervax induced 1477 upregulated and 1583 downregulated DEGs (Extended Data Fig. 2b). Among the mPervax-upregulated genes, a prominent cluster of innate immune mediators was observed including neutrophil chemotactic and activating factors (*Cxcl2, Cxcl3, Cxcl5*), type I interferons (*Ifna* family members and *Ifnb1*), and proinflammatory cytokines and myeloid growth factors (*Il6, Csf3*), where are closely associated with myeloid cell recruitment and IFN signaling activation (Extended Data Fig. 2b). To further dissect the functional programs underlying the transcriptional divergence between the two vaccines, all DEGs between mPervax and Boostrix groups were subjected to Gene Ontology (GO) and Kyoto Encyclopedia of Genes and Genomes (KEGG) pathway enrichment analyses. The enriched GO terms were overwhelmingly immune-related, including cytokine-mediated signaling and chemokine activity, leukocyte and neutrophil migration and chemotaxis, regulation of innate and inflammatory responses, I-κB kinase/NF-κB signaling, responses to type I interferon and lipopolysaccharide, macrophage activation, as well as antigen processing and presentation and MHC-related terms (Fig. 2c). In consistent, KEGG analysis highlighted cytokine-cytokine receptor interaction, chemokine signaling, Toll-like receptor and NOD-like receptor signaling, NF-κB and TNF signaling, IL-17 signaling, NK cell-mediated cytotoxicity, and antigen processing and presentation, which are highly involved in the establishment of anti-pertussis immunity (Fig. 2d).

**Figure 2.**
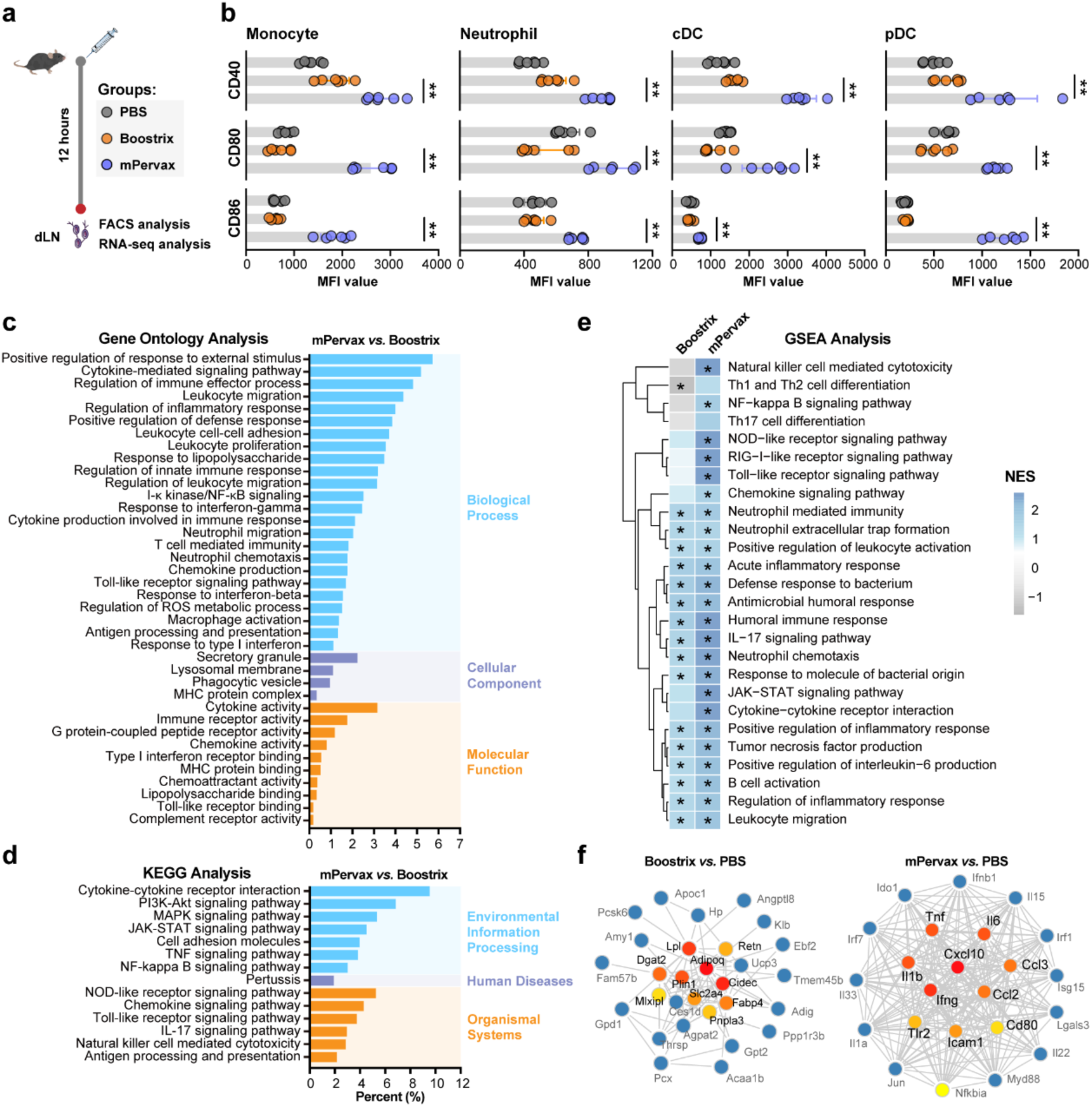
Comparative analyses of innate immune profiles by mPervax and Boostrix. **a,** Experimental design schematic. C57BL/6 mice (n=6/group) were i.m administered with mPervax (15 μg), Boostrix (1/3 human dose) or PBS. 12 hours post injection, dLNs were collected for analyses. **b,** Expression of CD40, CD80, CD83 on the indicated innate immune cell subsets was evaluated by flow cytometry. Mean fluorescence intensity (MFI) are shown. Data are presented as mean ± SEM. Unpaired Mann–Whitney U test was used for comparison. \*\**p* < 0.01. **c,** dLNs (4 biological replicates/group) were subjected to bulk RNA sequencing. Gene Ontology (GO) enrichment analysis of differentially expressed genes (DEGs) is shown. DEGs were identified using the criteria of *p* < 0.05 and |fold change| ≥ 2. **d,** Kyoto Encyclopedia of Genes and Genomes (KEGG) pathway enrichment analysis of DEGs is shown. **e,** Heatmap of gene set enrichment analysis (GSEA) results for modules related to immune responses. Normalized enrichment scores (NES) across treatment groups are displayed. \**p* < 0.05 indicates statistical comparison to the PBS group. **f,** Protein-protein interaction (PPI) network of DEGs constructed using data from the STRING database.

We further performed Gene Set Enrichment Analysis (GSEA) using gene lists from the Boostrix-PBS and mPervax-PBS comparisons, and the normalized enrichment scores (NES) were depicted as a heatmap (Fig. 2e). A large number of innate immune modules including pattern-recognition receptor signaling, chemokine signaling, neutrophil-mediated immunity and antibacterial defense response were upregulated upon both vaccination, which was more prominent in the mPervax group evidenced by higher NES. In addition, immune modules associated with NK cell-mediated cytotoxicity, Th1/Th2 and Th17 cell differentiation and NF-κB signaling were induced by mPervax yet unaltered or even decreased upon Boostrix vaccination, which suggested that these specific immune priming programs were not engaged by Boostrix at the early stage after vaccination. Consistent with these, protein-protein interaction (PPI) network analysis of the DEGs pinpointed the hub genes underlying each transcriptional program (Fig. 2f). The mPervax hubs were dominated by innate immune and inflammatory mediators, including the cytokines *Il1b, Il6, Tnf and Ifng*, the chemokines *Cxcl10 and Ccl2*, the pattern recognition receptor *Tlr2*, and the co-stimulatory molecule *Cd80*. In sharp contrast, the Boostrix hubs were mainly composed of lipid-metabolic regulators, including genes mediating lipid uptake, synthesis, and storage (*Lpl, Dgat2, Fabp4,* and *Plin1*) together with adipokine genes (*Adipoq* and *Retn*). Together, these data indicate that mPervax is strikingly more potent at activating innate immune compartment comprehensively in the vaccine injection site draining LNs at both cellular and transcriptional levels.

### mPervax induces robust Ab responses with enhanced magnitude and functionality than Boostrix

Immunogenicity of the different pertussis vaccine formulations were thoroughly evaluated in C57BL/6 mice upon a prime-boost vaccination regimen, with particular emphasis on the magnitude and function of Ab response, memory B cell (MBC) and T cell response (Fig. 3a). With regard to Ab response, two doses of mPervax vaccination induced significantly higher levels of anti-PT and anti-FHA IgG than Boostrix. Anti-PRN IgG titer also showed a clear trend of increase in the mPervax group (Fig. 3b). We also noticed that dual-component PT/FHA-mRNA-LNP vaccine elicited robust IgG against PT and FHA at comparable levels to that by mPervax, whereas mono-component PT-mRNA-LNP vaccine induced anti-PT IgG only (Fig. 3b), which verified that there was no inter-antigen interference among the antigen components within mPervax formulation. Further by dissecting Ab subclass composition, mPervax elicited both antigen-specific IgG1 and IgG2c responses, whereas Boostrix-induced IgG was exclusively restricted to IgG1 with a Th2 phenotype (Fig. 3c). The high magnitude of IgG2c elicited by mPervax across all three antigens highlighted a more Th1-prone immune response generated upon mRNA vaccination.

**Figure 3.**
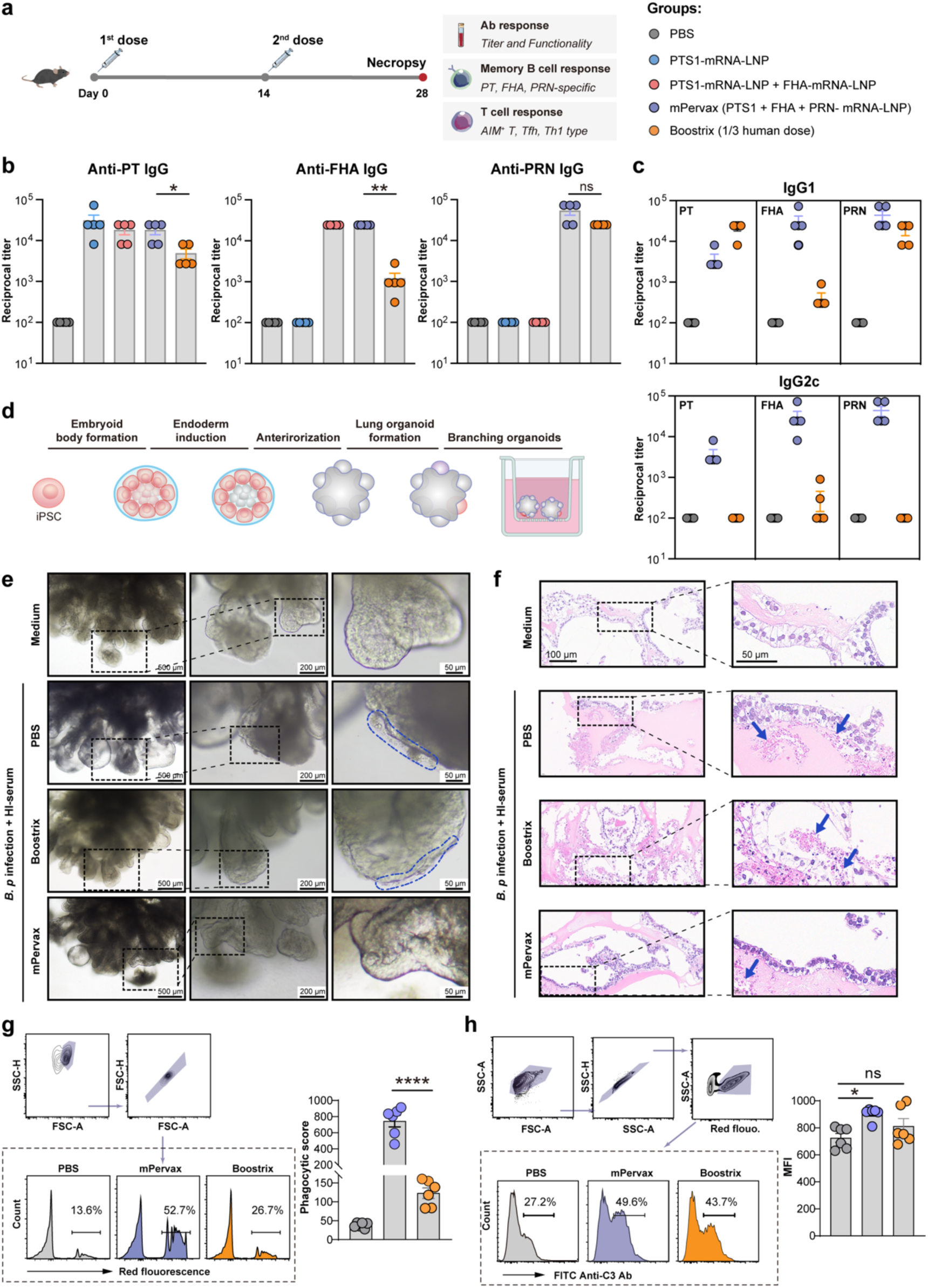
mPervax elicits robust Ab responses with higher magnitude and functionality than Boostrix. **a,** Experimental design schematic. C57BL/6 mice (n=5/group) were immunized with different vaccine formulations at an interval of 2 weeks. Antibody, memory B cell (MBC) and T cell responses were evaluated 2 weeks after the boost immunization. **b,** Serum IgG specific to PT, FHA and PRN were determined. Endpoint titers are shown. Data are presented as mean ± SEM. Unpaired Mann–Whitney U test was used for comparison. \**p* < 0.05, \*\**p* < 0.01. **c,** Antigen-specific IgG1 and IgG2 are shown. **d,** Experimental schematic for the establishment of human iPSC-derived branching lung organoid model. **e,** Heat-inactivated sera from immunized or PBS treated mice were pre-incubated with *B. pertussis* (strain BP18323) prior to infection of lung organoids. Morphological changes were examined. Representative images from two independent experiments are shown. **f,** Hematoxylin and eosin (H&E) staining of lung organoids. Arrows in blue color indicate histopathological changes. **g-h,** Ab-dependent cellular phagocytosis (ADCP) and Ab-dependent complement deposition (ADCD) activity of vaccine-induced Abs was assessed by flow cytometry (n=6). Data are presented as mean ± SEM. Mann–Whitney U test was used for comparison. \**p* < 0.05, \*\*\*\**p* < 0.0001.

We further evaluated the functions of vaccine-induced Abs using multiple assays. A human induced pluripotent stem cells (iPSC)-derived lung organoid model was established and deployed for assessing the capacity of Abs in blocking *B. pertussis* infection (Fig. 3d). In this regard, *B. pertussis* was pre-incubated with heat-inactivated sera collected from PBS-treated mice or vaccinated-mice prior to lung organoid infection. Compared to untreated lung organoid showing an intact epithelial surface, *B. pertussis* infection induced epithelial damage and reduced surface integrity (Fig. 3e). Presence of heat-inactivated sera from vaccinated mice could ameliorate cell damage and preserve epithelial integrity, which was more effective in mPervax group (Fig. 3e). The lung organoids were further subjected to histopathological assessment. In line with the morphological appearance, uninfected organoids maintained a well-preserved architecture characterized by a continuous monolayer of columnar epithelial cells enclosing a clear central lumen (Fig 3f). Upon bacterial challenge, organoids exhibited prominent focal disruption of the epithelial lining with irregular cell morphology and accumulation of cellular debris and eosinophilic material within the lumen. Notably, the lung organoids in culture supplemented with sera from mPervax-vaccinated mice were well preserved, which was characterized by a continuous epithelial lining and minimal intraluminal debris (Fig. 3f). These findings demonstrated that Abs elicited by mPervax were more potent at blocking *B. pertussis* infection to protect against tissue damage. While in contrast, Abs from Boostrix-immunized mice showed only partial protection. Beyond the neutralizing capacity, we also deployed systems serology approach to evaluate Fc-mediated Ab functions, with particular focus on the Ab-dependent cellular phagocytosis (ADCP) and Ab-dependent complement deposition (ADCD). mPervax-induced Abs exhibited strikingly higher ADCP activity than the Boostrix control (Fig. 3g), while the ADCD capacity of Abs was depicted at a comparable level between two groups (Fig. 3h). Together, these data show that mPervax elicits robust and high-quality Ab responses with enhanced magnitude and effector functions than Boostrix in mice, which could contribute to more efficient control of *B. pertussis* infection.

### mPervax elicits stronger CD4^+^ T cell responses than Boostrix

Effective protection against pertussis requires both specific Abs and CD4^+^ T cells^28^. It has been well-evidenced that Th1 and/or Th17-type CD4^+^ T cells are critical for controlling *B. pertussis* infection^36^. To characterize this, we first used an ELISpot assay to analyze the induction of splenic IFN-γ^+^ T cells specific to the three antigens or a pan-pertussis peptide pool. Two weeks after the boost, all animals in mPervax group showed strong induction of pertussis-specific T cells, in contrast to the Boostrix-immunized animals showing lower responses (Fig. 4a-b). We further used an intracellular cytokine recall assay to quantify the pertussis-specific CD4^+^ T cells producing IFN-γ, TNF or IL-17A upon peptide pool stimulation (Fig. 4c). Likewise, mPervax was able to induce a significantly higher frequency of IFN-γ^+^ or TNF^+^ CD4^+^ T cells than Boostrix. And a minimal level of IL-17A^+^ T cells were also induced upon vaccination in both two groups (Fig. 4d). We also analyzed the changes of T follicular helper (Tfh) cells that are critical for providing helper signals to B cells for geminal center reaction (Fig. 4e). In mPervax-immunized mice, splenic Tfh cells that are specific to PT and FHA showed a clear increase in frequency upon antigen stimulation. Notably, the expanded Tfh cell populations showed a more activated phenotype, in contrast to the Boostrix group showing a minimal level of cell activation (Fig. 4f). These data collectively indicate that mPervax is superior to Boostrix at inducing strong Th1-biased pertussis-specific CD4^+^ T and Tfh cell responses.

**Figure 4.**
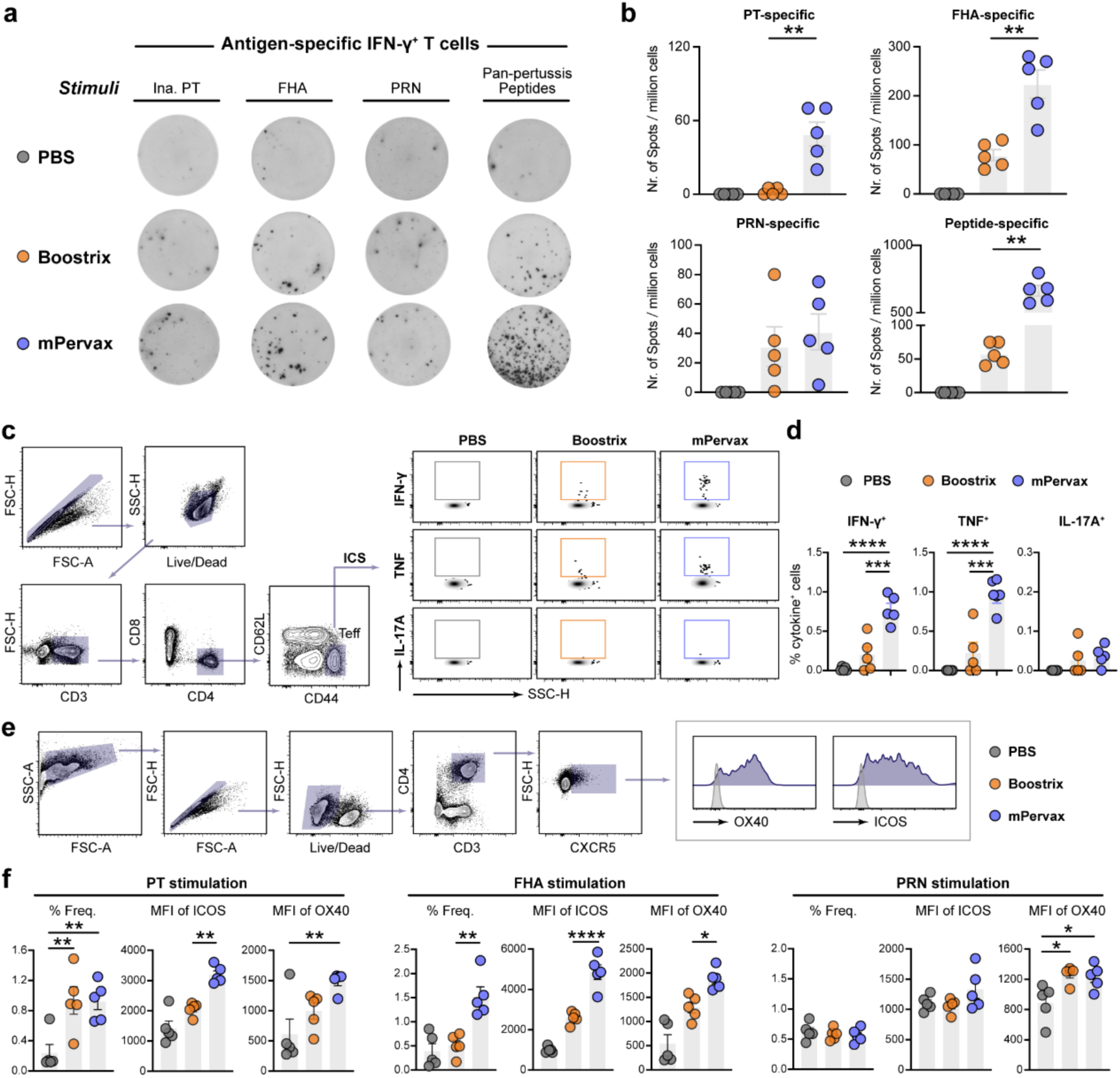
mPervax induces stronger Th1-type T cell and Tfh responses than Boostrix. C57BL/6 mice (n=5/group) were immunized with mPervax (15 μg), Boostrix (1/3 human dose) or PBS at an interval of 2 weeks. T cell response was evaluated at day 14 post the boost immunization. **a,** Splenocytes were stimulated with inactivated PT, FHA, or PRN (each at 2 μg/mL) or a pertussis peptide pool (100 ng/mL) for 20 hours. Frequencies of IFN-γ producing T cells were measured by ELISpot assay. Data from representative animals are shown. **b,** Numbers of antigen-specific T cells were enumerated and indicated as spots per million stimulated cells. Data are presented as mean ± SEM. Mann–Whitney U test was used for comparison. \*\**p* < 0.01. **c,** Frequencies of cytokine-producing effector CD4^+^ T cells upon pertussis peptides stimulation were evaluated by flow cytometry. Gating strategy is shown. **d,** Quantification of IFN-γ, TNF or IL-17A-producing CD4^+^ T cells. Negative control values from unstimulated cells were subtracted. One-way analysis of variance (ANOVA) with Tukey’s post hoc test was used for statistical comparison. Data are presented as mean ± SEM. \*\*\**p* < 0.001, \*\*\*\**p* < 0.0001. **e,** Gating strategy for Tfh cell population (CD3^+^CD4^+^CXCR5^+^) and the surface expression of OX40 and ICOS. **f,** Splenocytes were stimulated with inactivated PT, FHA or PRN (each at 2 μg/mL) for 18 hours. Frequencies of Tfh cells and their surface expression levels of ICOS and OX40 are shown. One-way analysis of variance (ANOVA) with Tukey’s post hoc test was used for statistical comparison. Data are presented as mean ± SEM. \**p* < 0.05, \*\**p* < 0.01, \*\*\*\**p* < 0.0001.

### mPervax induces efficient protection against *B. pertussis* challenge in adult mice and confers passive protection to neonatal mice upon maternal immunization

To evaluate the protective efficacy of each vaccine, we administered mice with two doses of pertussis vaccines followed by intranasal challenge with 5 × 10^8^ CFU of virulent *B. pertussis* (WHO reference strain 18323) and evaluation of pulmonary bacterial burden and lung tissue damage (Fig. 5a). The mono-component PT-mRNA-LNP vaccine and dual-component PT/FHA-mRNA-LNP vaccine were also assessed for their protective capacity in parallel. To ensure successful immunization, we measured Ab response one day prior to challenge and found that all vaccinated mice demonstrated robust levels of anti-PT IgG (Extended Data Fig. 3). At day 7 post challenge, unvaccinated mice exhibited high bacterial loads in the lungs, while the PT-mRNA-LNP and PT/FHA-mRNA-LNP vaccines provided low to moderate levels of protection in terms of reduction of bacterial burden (Fig. 5b). Notably, mPervax vaccination induced a highly efficient clearance of bacteria, with 4/6 animals showing sterilizing immunity with undetectable pulmonary bacterial loads. By contrast, Boostrix vaccination reduced bacterial load at a moderate level, but its efficacy was less robust than mPervax (Fig. 5b). We further performed hematoxylin and eosin (H&E) staining of lung tissues for histopathological assessment. Unvaccinated mice that were challenged with *B. pertussis* revealed severe pulmonary damage, which was characterized by thickening of alveolar wall, inflammatory cell infiltration around the bronchi and alveoli, as well as pulmonary edema (Fig. 5c). In comparison, the severity of pulmonary lesions was significantly attenuated in both two vaccine groups showing less inflammatory cell infiltration and well-preserved alveolar architecture (Fig. 5c, Extended Data Fig. 4a).

**Figure 5.**
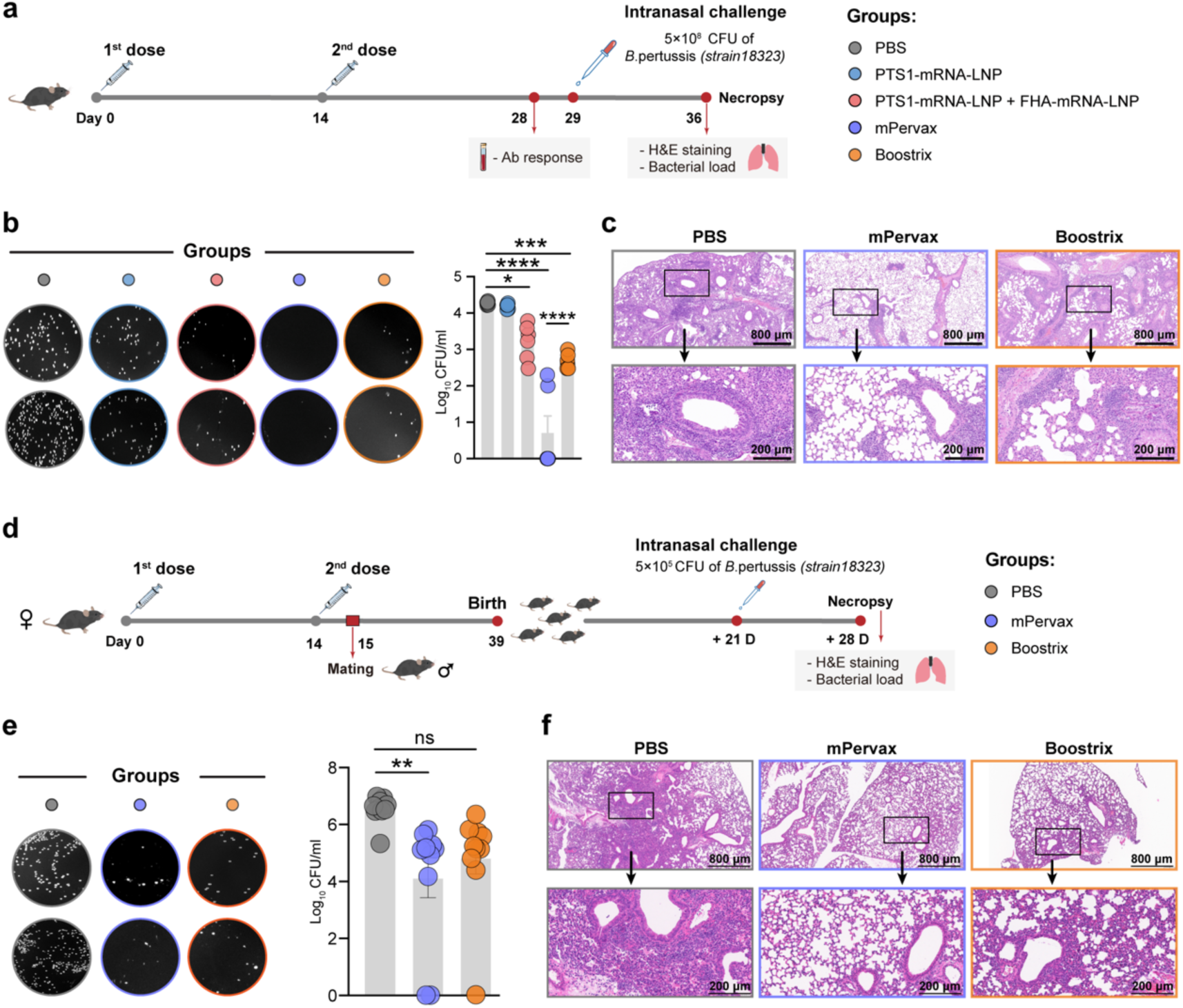
mPervax demonstrates higher protective efficacy against *B. pertussis* challenge than Boostrix. **a,** Experimental design schematic. C57BL/6 mice (n=6/group) were immunized with different vaccine formulations at day 0 and day 14. Serum samples were analyzed for Ab response 2 weeks after boost immunization. At day 29, mice were intranasally challenged with 5 × 10⁸ CFU of *Bordetella pertussis* strain 18323. Seven days post bacterial challenge, animals were necropsied for analyses. **b,** Bacterial burdens were examined and shown as Log_10_ (CFU) per mL homogenate of left lungs. Data are presented as mean ± SEM. One-way analysis of variance (ANOVA) with Tukey’s post hoc test was used for statistical comparison. \**p* < 0.05, \*\*\*\**p* < 0.0001. **c,** Representative images of H&E staining of the right lung tissues. **d,** Female mice immunized with two doses of pertussis vaccines at day 0 and day 14 were subjected to mating on day 15. Neonatal mice (n=10/group) at the age of 3 weeks were intranasally challenged with 5 × 10^5^ CFU of *B. pertussis*. **e,** Bacterial burdens in left lungs were examined and shown as Log_10_ (CFU) per mL of lung homogenate. Data are presented as mean ± SEM. One-way analysis of variance (ANOVA) with Tukey’s post hoc test was used for statistical comparison. \*\**p* < 0.01. **f,** Representative images of H&E staining of the right lung tissues.

Maternal pertussis vaccination has been implemented in several countries, which provides significant health benefits by decreasing pertussis morbidity and mortality in infants ^37,38^. In this study, we evaluated whether maternal antibodies induced by mPervax or Boostrix vaccination could protect newborn mice against *B. pertussis* challenge. In this context, female mice that were primarily immunized with two doses of pertussis vaccines at day 0 and day 14 were subjected to mating on day 15, and were allowed to give birth naturally on day 39. The neonatal mice were subsequently challenged with 5 × 10^5^ CFU of *B. pertussis* intranasally three weeks post birth and evaluated for protection (Fig. 5d). As expected, the bacterial burden in the lungs of offsprings born to unvaccinated mice remained at a high level. Maternal vaccination largely protected the pups against pulmonary bacterial colonization, which was significantly more effective upon mPervax vaccination (Fig. 5e). In addition, pups born to the mPervax-vaccinated mice showed relatively lower bacterial burdens in the lungs than the benchmark vaccine group (Fig. 5e). Histopathological analysis of the neonatal lungs demonstrated that maternal pertussis vaccination effectively prevented *B. pertussis*-induced pulmonary injury in neonates, with no difference observed between the two vaccine groups (Fig. 5f, Extended Data Fig. 4b). We further measured the levels of representative inflammatory cytokines in sera collected from pups 7 days post bacterial challenge. Relative to the mice in PBS group, pups born to mPervax- or Boostrix-immunized mice had significantly lower serum levels of IL-6, MCP-1 and IFN-γ upon infection (Extended Data Fig. 5). Together, these data indicate that mPervax provides robust protection against *B. pertussis* challenge in adult mice, and maternal mPervax vaccination confers efficient passive protection to neonatal offspring.

### mPervax confers superior protection against antibiotic-resistant *B. pertussis* variants than Boostrix

Emerging *B. pertussis* isolates harboring genetic mutation, antigen deficiency, and antimicrobial resistance have increasingly been reported worldwide, which is potentially attributed to vaccine-driven selective pressure ^39,40^. In this study, the wide-type reference strain BP18323 and three unique drug-resistant clinical variants BP18216, BRK03 and BRK15 were obtained and well characterized (Fig. 6a). BRK03 strain is deficient in PRN, and BRK15 strain is a recently emerging variant deficient in both PRN and FHA^17^. These representative strains were first examined for their antimicrobial susceptibility using disk diffusion assay, which revealed that BP18216, BRK03 and BRK15 variants exhibited strong resistance to azithromycin (AZM), erythromycin (ERY) and clindamycin (CD), while in contrast the BP18323 strain was highly susceptible (Fig. 6b). Furthermore, minimum inhibitory concentration (MIC) testing against a broader panel of antibiotics showed that BP18216, BRK03 and BRK15 variants are highly resistant to AZM, ERY, CD and clarithromycin with determined MIC values above 256 mg/L (Fig. 6c).

**Figure 6.**
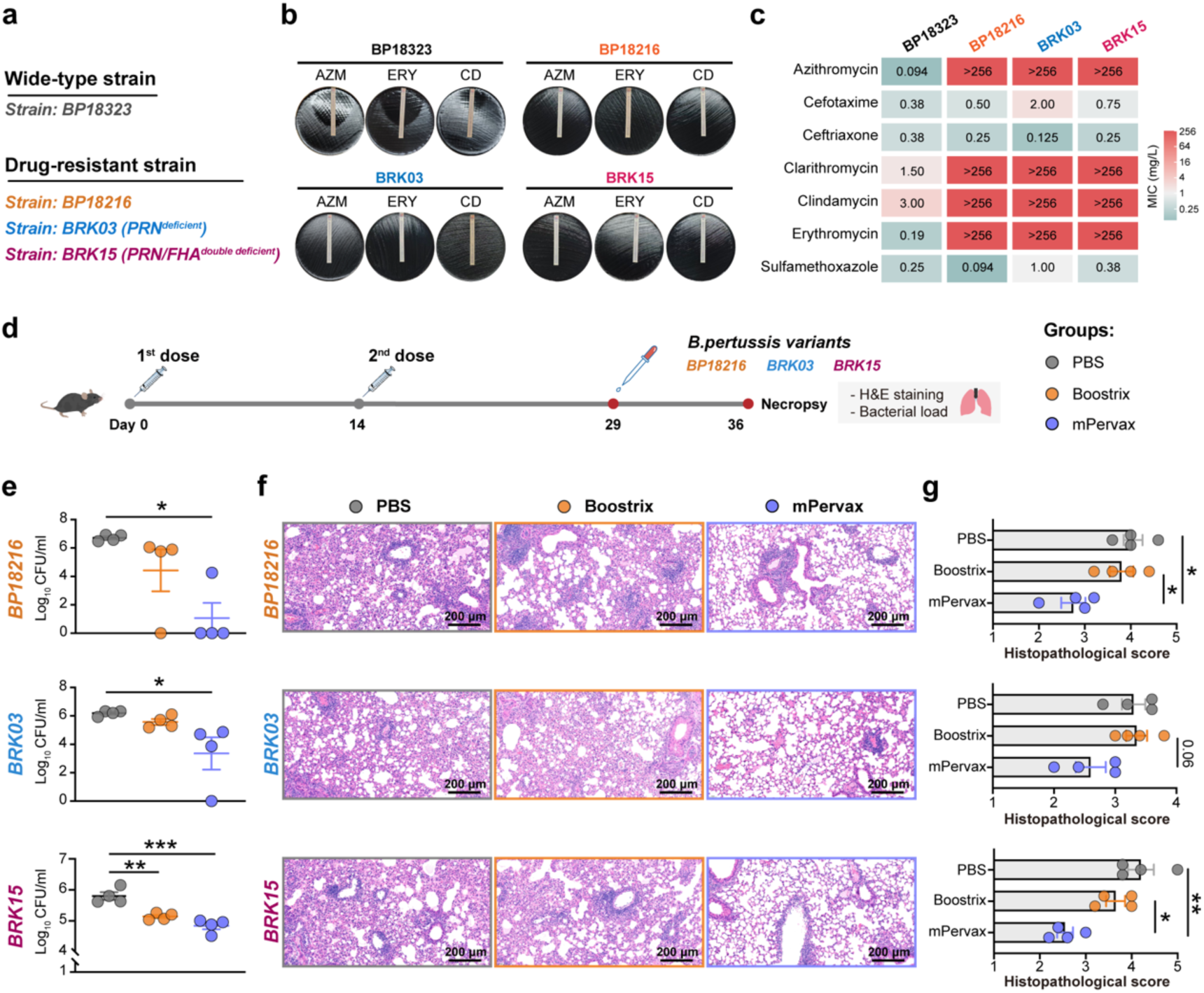
mPervax confers superior protection against antibiotic-resistant *B. pertussis* variants than Boostrix. **a,** Information of representative *B. pertussis* strains. **b,** Susceptibility of the four *B. pertussis* isolates to AZM, ERY and CD was evaluated. Representative images of disk diffusion assay are shown. **c,** Minimum inhibitory concentration (MIC) testing of the indicated antibiotics against the four *B. pertussis* isolates. **d,** Experimental design schematic. C57BL/6 mice (n=4/group) were immunized with mPervax (15 μg) or Boostrix (1/3 human dose) at day 0 and day 14, followed by intranasal challenge with antibiotic-resistant variants. **e,** Bacterial burdens in left lungs were examined and shown as Log_10_ (CFU) per mL of lung homogenate. Data are presented as mean ± SEM. One-way analysis of variance (ANOVA) with Tukey’s post hoc test was used for statistical comparison. \**p* < 0.05, \*\**p* < 0.01, \*\*\**p* < 0.001. **f,** Representative images of H&E staining of the right lung tissues. **g,** Histopathological scores of the H&E staining. Data are presented as mean ± SEM. One-way analysis of variance (ANOVA) with Tukey’s post hoc test was used for statistical comparison. \**p* < 0.05, \*\**p* < 0.01.

We next compared the protective capacity of mPervax and Boostrix against antibiotic-resistant variants using the murine challenge model (Fig. 6d). At day 7 post challenge, mice from the PBS group exhibited high levels of bacterial burdens in the lungs across all three challenge groups. While Boostrix vaccination reduced bacterial loads to a certain extent, mice vaccinated with mPervax were better protected from pulmonary bacterial colonization. This protective efficacy was more pronounced against the BP18216 and BRK03 variants, which showed a potent reduction in bacterial loads compared to the Boostrix group (Fig. 6e). Histopathological evaluation further revealed a significant protection against lung damage in mPervax-vaccinated group. Interestingly, although Boostrix vaccination showed a moderate ability in reducing bacterial loads, the vaccinated mice exhibited a very limited mitigation of pulmonary inflammation and damage compared to the PBS and mPervax groups (Fig. 6f-g). Collectively, these findings indicate that mPervax provides robust and superior protection against emerging antibiotic-resistant *B. pertussis* variants, which outperforms the licensed Boostrix vaccine in reducing both bacterial burden and lung injury.

### Heterologous prime-boost with mPervax induces higher vaccine responses than homologous prime-boost

Pertussis vaccines are conventionally administered as a primary series in infants and children, followed by booster doses for adolescents or adults. Given this established pediatric vaccination schedule, we sought to evaluate the potential of mPervax as a booster dose following a primary Boostrix vaccination regimen. To this end, mice were primed with two doses of Boostrix at day 0 and day 14 followed by either a heterologous booster with mPervax or a homologous booster with Boostrix on day 49 (Fig. 7a). Serum IgG specific to PT, FHA and PRN were monitored longitudinally. We found that the heterologous mPervax booster elicited higher titers of antigen-specific IgG than Boostrix booster, which was more prominent for anti-FHA and anti-PRN IgG (Fig. 7b). In addition, the mPervax boost vaccination induced a significantly higher level of anti-PT IgG2c and IgG2c/IgG1 ratio (Fig. 7c), which indicates that the mRNA vaccine booster effectively promoted a shift towards a Th1-biased humoral response. We also assessed antigen-specific T cell responses by stimulating splenocytes with a pan-pertussis peptide pool and measured the frequency of IFN-γ–producing CD4⁺ T cells. Compared to the homologous boost group, the heterologous mPervax booster induced a higher Th1-type cellular immunity (Fig. 7d). These data collectively indicate that a heterologous booster strategy using mPervax not only promotes vaccine responses but also shift the early established immunity towards a Th1 phenotype, thereby overcoming the Th2-skewing limitation of conventional aPV.

**Figure 7.**
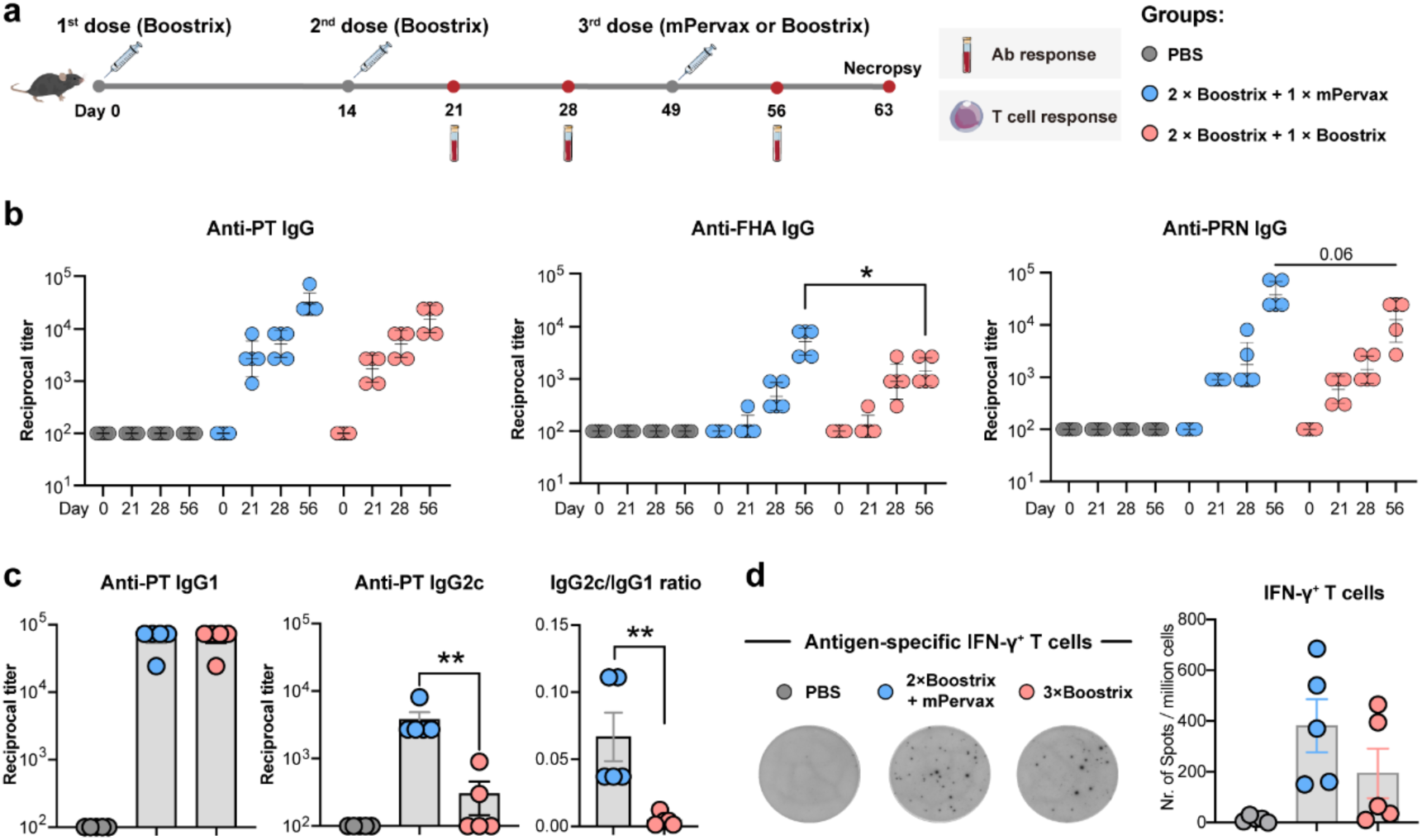
Heterologous mPervax boost vaccination promoted Th1-type anti-pertussis immunity. **a,** Experimental design schematic. C57BL/6 mice (n=5/group) were primarily immunized with two doses of Boostrix (1/3 human dose) at day 0 and day 14, followed by a heterologous booster with mPervax (15 μg) or a homologous booster with Boostrix (1/3 human dose) at day 49. Serum and spleen samples were collected at the indicated time points for analyses. **b,** Serum IgG specific to PT, FHA and PRN were determined. Endpoint titers are shown. Data are presented as mean ± SEM. Mann–Whitney U test was used for comparison. \**p* < 0.05. **c,** Anti-PT IgG1, IgG2c titers and IgG2c/IgG1 ratio at day 63 are shown. Data are presented as mean ± SEM. Mann–Whitney U test was used for comparison. \*\**p* < 0.01. **d,** At the day of necropsy, pertussis-specific T cell response upon peptides pool stimulation was evaluated by ELISpot assay. Data from representative animals are shown. Numbers of antigen-specific IFN-γ-secreting T cells were enumerated and indicated as spots per million stimulated cells. Data are presented as mean ± SEM.

## Discussion

The resurgence of pertussis represents a critical and unresolved public health challenge and the widespread use of aPVs has fundamentally shaped the global epidemiological landscape^41^. While aPVs have a more favorable safety profile than wPVs, substantial evidence indicates that this type of vaccine predominantly elicits a Th2-skewed vaccine response that is suboptimal for controlling *B. pertussis* infection^8,9^. Moreover, vaccine and antibiotic-driven selection pressure has accelerated the emergence of antigen-deficient and macrolide-resistant clinical variants, which further complicates pertussis prevention and clinical management^18^. In this study, we rationally engineered a tricomponent mRNA pertussis vaccine (mPervax) and demonstrated its comprehensive superiority over the licensed aPV Boostrix and its unique capacity to shape pulmonary immune microenvironment favoring efficient protection against *B. pertussis*.

A fundamental limitation of conventional aPVs is their restricted ability to stimulate comprehensive and functional anti-pertussis immunity. Our data reveal that mPervax not only orchestrates a broader and more potent early innate immune activation but also elicits high-magnitude and high-quality Ab and cellular responses. These further translate into superior protective capacity against both wild-type and antibiotic-resistant variants. Regarding the early response, transcriptomic and flow cytometric profiling showed that mPervax triggered a remarkably stronger innate activation in draining LNs than Boostrix, which involves coordinated triggering of multiple pattern-recognition receptors including Toll-like receptor, NOD-like receptor, RIG-I-like receptor and their downstream signaling pathways leading to cell activation and migration. These findings suggest that mRNA vaccines formulated with ionizable lipid LNP are unique in their capacity to activate innate compartment compared to other vaccine types, which is mediated by the internal adjuvant property of both mRNA payload and lipid carrier^31,42,43^.

Vaccine-elicited innate signature fundamentally determines efficient priming of adaptive immunity. Indeed, mPervax drove a profound Th1-polarized CD4^+^ T cell response that was characterized by high frequencies of IFN-γ and TNF-producing cells against all three vaccine antigens. This Th1 bias is mechanistically beneficial for controlling *B. pertussis* infection since IFN-γ secreted by Th1-type T cells can potently activate innate effector cells, such as neutrophils and macrophages, to enhance their phagocytic and bactericidal activity. While in stark contrast, Boostrix vaccination elicited a minimal level of Th1 response, which is also consistent with other previous studies^11,44,45^. Moreover, we found that a heterologous mPervax boost upon primary Boostrix immunization could shift the imprinted T cell immunity towards a protective Th1 phenotype. So far, limited evidence is available on whether an mRNA vaccine could reshape the immune profile established by prior aPV vaccination. Our data suggests that mPervax may serve not only as a standalone next-generation pertussis vaccine but also hold promise as a powerful booster to optimize the immune profile in populations already primed with conventional aPVs.

Beyond T cell immunity, Ab responses are also critical in terms of both magnitude and functionality. Our data indicate that mPervax not only elicited higher antigen-specific IgG titers than Boostrix, but also generated Abs with enhanced functional activity. In a physiologically relevant lung organoid infection model, sera from mPervax-immunized mice exhibited superior neutralizing capacity in preserving epithelial integrity and preventing tissue damage upon infection. The enhanced functionality of Abs is further reflected by higher ADCP activity that is crucial for eliminating pathogen through engagement with innate effector cells through opsonization. Mechanistically, the highly activated antigen-specific Tfh cells induced by mPervax likely provide strong helper signals to B cells within germinal centers, which drives the class-switching towards Th1-associated IgG2c accompanied by affinity maturation. Likewise, in the homologous or heterologous prime-boost experiment, a booster dose of mPervax upon Boostrix immunization promoted anti-pertussis Ab responses more robustly than the homologous Boostrix boost, and the generated Abs were more shifted towards a Th1 phenotype. This finding is closely relevant to adult pertussis booster strategy that is being employed worldwide since improving the durability and quality of vaccine-induced Ab response may help reduce household transmission to infants^46^. Altogether, mPervax-elicited Abs are not merely abundant, but highly functional and optimal for effective bacterial clearance.

In addition to immunogenicity evaluation, the protective potential of mPervax and Boostrix was thoroughly examined and compared in a well-established murine challenge model. Currently, macrolide-resistant *B. pertussis* (MRBP) isolates have emerged and are posing significant challenges for pertussis control^47^. High prevalence of MRBP among clinical isolates has been reported worldwide, which is particularly serious in China with an estimated rate > 50%^18^. Moreover, macrolide treatment failure in MRBP infection has been frequently documented, which underscores direct threat that MRBP poses to clinical treatment^48,49^. In our study, mPervax vaccination provided robust protection against both WHO reference strain 18323 and critically, the representative MRBP variants including strain BP18216, BRK03 (PRN ^deficient^) and BRK15 (PRN/FHA ^double-deficient^). In contrast, protective efficacy of Boostrix against all the four *B. pertussis* strains was notably reduced. This compromised efficacy against variants deficient in vaccine antigen is quite expected since traditional aPV relies heavily on Ab response against infection. However, we found that mPervax that encodes three principal antigens similar to Boostrix components retained an efficient protective capacity against antigen-deficient MRBP variants. This implies that protection induced by mPervax is not exclusively relied on FHA- or PRN-specific immunity. The underlying mechanistic advantage in this context is likely multifaceted that involves a well-coordinated operation between PT-specific Abs, strong Th1-type T cells and enhanced Fc-mediated effector functions engaging innate effector cells, which are independently of PRN and FHA recognition. Lastly, the clinical potential and significance of mPervax further extends to the scenario of maternal immunization. We showed that maternal mPervax vaccination conferred highly efficient passive protection to neonatal mice, which significantly reduced pulmonary bacterial burdens and tissue damage upon bacterial challenge. This highlights the potential of mPervax as a maternal vaccine to bridge the immunity gap before infants receive their primary pertussis vaccination.

Our study has some limitations. More in-depth investigations into the mode of action of mPervax in its ability to alter the local respiratory immune environment when administered via non-systemic route remains largely unknown. It would be particularly interesting to evaluate the ability of mPervax in preventing nasopharyngeal colonization and transmission in suitable animal models. Future studies should also assess the long-term durability of the protective immunity induced by mPervax and the safety profile awaits thorough assessment for future potential translation to human studies. Despite these limitations, this study reports on the rational development of a highly effective tricomponent mRNA pertussis vaccine candidate, which demonstrated superior immunogenicity and protective capacity over licensed conventional aPV against emerging antibiotic-resistant variants. Our findings provide compelling evidence showing that mRNA platform is a highly promising avenue for next-generation pertussis vaccine development. By activating well-coordinated immune compartments across strong innate activation, robust Th1/Tfh cellular immunity, high-quality antibodies with enhanced effector functions and a favorable mucosal immune microenvironment beneficial for bacterial clearance, mPervax outperforms the licensed aPV in multiple preclinical contexts. This vaccine particularly addresses the key challenge of the emergence of antibiotic-resistant, antigen-deficient *B. pertussis* variants, paving the way for a novel intervention strategy to combat the global pertussis resurgence.

## Methods

### Ethics, animals, and immunization

Animal experiments were conducted in accordance with the Guidelines for the Care and Use of Laboratory Animals and the Ethical Committee of China Pharmaceutical University (approval no. CPUTML-LA-20240301 and YSL-202506036). C57BL/6 mice (6 weeks old) were purchased from Zhejiang Vital-River Laboratory Animal Technology Co., Ltd. and randomly assigned to the experimental groups. Animals were housed either in a specific pathogen-free (SPF) facility or in an animal biosafety level 2 (ABSL-2) facility where the temperature was maintained at 22 ± 1°C and a 12-hour light/dark cycle was used. For experiments evaluating vaccine immunogenicity and protective efficacy, mice were vaccinated intramuscularly (i.m.) following a prime-boost regimen at a 2-week interval. Boostrix vaccine was obtained from GlaxoSmithKline. mRNA vaccines were prepared in house. Serum samples were collected at the indicated timepoints. At the day of necropsy, mice were euthanized, and injection site draining lymph nodes or spleens were harvested and processed into single-cell suspensions for further analyses.

### mRNA design and synthesis

mRNAs encoding for vaccine antigens including PTS1-mRNA, FHA-mRNA and PRN-mRNA were designed and codon-optimized using proprietary algorithm mainly based on codon-adaptation index (CAI), minimum free energy (MFE) and translation initiation efficiency (TIE). All mRNA sequences were engineered with the 5ʹ-UTR of human α-globin, AES-mtRNR1 3ʹ-UTR and a 120 nt poly A tail, and the sequences were indicated in Extended Data Table 1. mRNAs with high purity above 95% were synthesized using a well-established T7 RNA polymerase-mediated in vitro transcription (IVT) protocol as previously reported by our group^20,31,32^. During the IVT procedure, mRNAs were modified with N1-methylpseudouridine (NEB) and co-capped with Cap 1 analogue (CleanCap AG, C_32_H_43_N_15_O_24_P_4_, TriLink). Following this step, mRNAs were purified by LiCl precipitation and resuspended in DNase/RNase-free TE buffer.

### LNP formulation and characterization

mRNA vaccine was formulated into LNP using a well-established protocol as described in our previous studies^21,22,50^. In brief, an organic phase containing four lipid components dissolved in ethanol at a molar ratio of 50:10:38.5:1.5 (ionizable lipid SM102: DSPC: cholesterol: DMG-PEG2000, all purchased from SINPOEC) was mixed with an aqueous phase containing mRNAs at a volume ratio of 3: 1 using an INano™ L microfluidic instrument (INano^L^ from Micro&Nano Biologics) at a total flow rate of 12 mL/min. LNP formulation was subsequently diluted with PBS and ultrafiltered using 50-kDa Amicon ultracentrifugal filters (Merck-Millipore). Particle size, polydispersity index (PDI) and zeta potential of the formulation were measured using a NanoBrook Omni Zeta-Plus instrument (Brookhaven Instruments). Encapsulation efficiency and mRNA concentration were measured using the Quant-iT™ RiboGreen RNA assay (Invitrogen, US).

### Measurement of antigen-specific Abs

PT, FHA and PRN antigens (gifted by National Institutes for Drug Control, NIDC) were coated onto 96-well plates (Greiner Bio-One) at 100ng/well and incubated overnight at 4° C. Plates were washed 3 times with PBS containing 0.075% Tween 20 (PBST) and blocked with 2% bovine serum albumin (BSA) for 2 hours at room temperature (RT). For detection of antigen-specific IgG, IgG1 and IgG2c, serially diluted serum samples (starting from 1: 100) were added into the wells and incubated for 2 hours at 30 °C. Then, HRP-conjugated goat anti-mouse IgG (1:50,000 dilution; Abcam), IgG1 (1:5,000 dilution; SouthernBiotech), or IgG2c (1:5,000 dilution; SouthernBiotech) antibodies were added and incubated for 1 hour at 30°C. Following washing steps, 3,3’,5,5’-Tetramethylbenzidine (TMB) substrate was used for development and the absorbance was measured at 450 nm using SpectraMax Absorbance Reader (Molecular Devices). Endpoint titers were defined as the reciprocal serum dilution that exceed 2.1-folded optical density (OD) value of the background.

### Antigen recall T cell assay

For the evaluation of antigen-specific T cell response, two million murine splenocytes were plated into 96-well U-bottom plates and stimulated with PepMix™ Pan-Pertussis peptide pool (10μg/mL, JPT Peptide Technologies) in the presence of brefeldin A (BioLegend) for 8 h at 37 °C. Cells cultured in the medium were used as negative controls. Cytokine production of antigen-specific T cells was evaluated by surface and intracellular staining (ICS) using Fixation/Permeabilization Solution Kit (BD Biosciences). Frequencies of cytokine-producing T cells were determined by flow cytometer. Background cytokine staining was subtracted, which was defined by staining in the samples incubated with medium alone. Data were acquired on BD Fortessa (BD Biosciences) and were analyzed using FlowJo V.10.1 (Tree Star). Antibodies used for flow cytometric analyses are listed in Extended Data Table 2. In some experiment, an Enzyme-linked immunospot (ELISPOT) assay was used for examining T cell response. In this regard, murine splenocytes (0.2 million cells /well) were incubated with or without heat-inactivated PT, FHA or PRN (2μg/mL) or with PepMix™ Pan-Pertussis peptide pool (10μg/mL, JPT Peptide Technologies) for 24 hours. Unstimulated cells served as negative controls. Frequency of IFN-γ–secreting T cells was determined using an IFN-γ ELISpot kit (Mabtech) according to the instructions. Spots were developed with BCIP/NBT substrate (Mabtech) and quantified using a CTL ImmunoSpot S6 Analyzer. Results are depicted as spot-forming cells (SFCs) per million stimulated cells.

### *B. pertussis* variants and antimicrobial susceptibility testing

*B. pertussis* isolates including the WHO reference strain BP18323 and macrolide-resistant variants BP18216, BRK03 and BRK15 were obtained from Chinese Center for Disease Control and Prevention. *B. pertussis* isolates were cultured on charcoal agar plates (CM119B, Oxoid, UK) supplemented with 10% (v/v) defibrinated sheep blood (Y099, Sanmu Animal Husbandry, China) at 37°C for 3–5 days. Detailed genomic sequences of BP18216, BRK03 and BRK15 variants have been uploaded to National Library of Medicine (NLM), National Center for Biotechnology Information (NCBI) at the National Institutes of Health (Submission ID: SUB16384001), which are available under BioProject: PRJNA1507474. Susceptibility of *B. pertussis* isolates to antibiotics was assessed and minimum inhibitory concentrations (MICs) of azithromycin, cefotaxime, ceftriaxone, clarithromycin, clindamycin, erythromycin, and sulfamethoxazole were determined using Liofilchem® MIC Test Strips (Italy) according to the manufacturer’s instruction. MICs were determined after 4 days of incubation. *Staphylococcus aureus* (ATCC 29213) and *Escherichia coli* (ATCC 25922) were used as the quality control strains. As the American Clinical and Laboratory Standards Institute (CLSI) and the European Committee on Antimicrobial Susceptibility Testing (EUCAST) do not establish breakpoints for *B. pertussis*, susceptibility results are presented as MIC values.

### *B. pertussis* murine challenge model

*B. pertussis* challenge experiments were performed in an animal biosafety level 2 (ABSL-2) facility. Unvaccinated and vaccinated C57BL/6 mice were anesthetized using 2% isoflurane and then intranasally challenged with 5 × 10⁸ CFU of different *B.* pertussis isolates in a volume of 50µL. In some experiment, C57BL/6 mice were intranasally challenged with 5 × 10^5^ CFU of *B. pertussis*. At day 7 post bacterial challenge, mice were euthanized by cervical dislocation under isoflurane anesthesia. The left lung tissues were harvested, homogenized and centrifuged at 300 × g for 10 min, and the supernatant was used for quantification of bacterial burdens. The right lung tissues were fixed in 4% paraformaldehyde (PFA), paraffin-embedded, sectioned, and hematoxylin and eosin (H&E) stained for histopathology evaluation.

### *B. pertussis* infection of lung organoid model

Branching lung organoid (BLO) model was established by differentiating human induced pluripotent stem cells (iPSCs) using STEMdiff™ Branching Lung Organoid Kit (STEMCELL Technologies) via a four-stage protocol according to the manual. At stage 1 (d0-d3), iPSCs were first induced to form definitive endoderm (DE). At stage 2 (d3-d6), the cells were differentiated into anterior foregut endoderm (AFE). At Stage 3 (d6-d14), AFE buds were cultured in suspension and further developed into lung bud organoids. At Stage 4 (after d14), the organoids were embedded in a Matrigel^®^ sandwich in Transwell^®^ inserts and cultured to form BLOs with branching airway structures. For the infection of BLOs, *B. pertussis* was used at an infection dose of MOI = 10, corresponding to approximately 1 × 10^4^ CFU. To test the blocking effect of sera collected from immunized mice, *B. pertussis* was incubated with heat-inactivated sera (56°C for 30 minutes) at 37°C for 1 hour to allow antibody binding to bacterial surface. Afterwards, bacteria-serum mixture was added to the organoid culture for incubation. Four hours later, the culture medium was refreshed, and the organoids were cultured for additional 48 hours. Organoids were assessed by bright-field microscopy for gross morphological changes and further processed to H&E staining for histopathological examination.

## Statistical analysis

Statistical calculations were performed using GraphPad Prism v6.0. Comparisons between groups were determined using one-way ANOVA with Tukey’s post hoc test or unpaired Mann–Whitney U test. A *p* value less than 0.05 was considered statistically significant.

## Acknowledgements

This work was supported by the National Key Research and Development Program of China (Grant No. 2025YFC2311900), Innovation and Entrepreneurship (ShuangChuang) Program of Jiangsu Province (Grant no. JSSCTD202447), Jiangsu Provincial Graduate Research and Practical Innovation Program (Grant no. KYCX25-1083). We thank the Targeted Discovery Center of China Pharmaceutical University for instrumental support on this study.

## Author contributions

Study design: A.L. and Z.S. Supervision and funding acquisition: A.L., Z.S., L.W. and M.Z. Investigation, methodology and validation: W.Z., J.C., G.W., J.W., S.Q., Z.H., H.D., J.M., J.M., X.S., K.G., P.X., Z.D., Q.A., S.Z., L.W. and A.L. Discussion: A.L., Z.S., L.W., W.Z., J.C., G.W., S.Z., X.G., Y.Y., X.M., L.Y. Writing, review and editing: A.L., W.Z. and G.W. All authors have read and approved the manuscript.

## Data availability

All data are available upon reasonable request to the corresponding author. Raw sequencing data related to the work on innate immune profiling have been deposited in the open OMIX platform the National Genomics Data Center (https://ngdc.cncb.ac.cn/omix; project no. PRJCA080928; Accession no.: CRA054879).

## Reference

1 GBD 2023 Pertussis Collaborators, Mosser, J. F. & Liu, J. Global, regional and national burden of pertussis in 1990-2023: a Global Burden of Disease study. Nat. Med. 32, 4021–4035 (2026).

2 Locht, C. Pertussis before, during and after COVID-19. EMBO Mol. Med. 17, 594–598 (2025).

3 World Health Organization. Immunization data. https://immunizationdata.who.int/ (2026).

4 Wang, S., Zhang, S. & Liu, J. Resurgence of pertussis: Epidemiological trends, contributing factors, challenges, and recommendations for vaccination and surveillance. Hum. Vaccin. Immunother. 21, 2513729 (2025).

5 Locht, C. Pertussis: acellular, whole-cell, new vaccines, what to choose? Expert Rev. Vaccines 15, 671–673 (2016).

6 Lin, A. et al. Live attenuated pertussis vaccine BPZE1 induces a broad antibody response in humans. J. Clin. Invest. 130, 2332–2346 (2020).

7 Leef, M., Elkins, K. L., Barbic, J. & Shahin, R. D. Protective immunity to *Bordetella pertussis* requires both B cells and CD4(+) T cells for key functions other than specific antibody production. J. Exp. Med. 191, 1841–1852 (2000).

8 Diavatopoulos, D. A. & Edwards, K. M. What Is Wrong with Pertussis Vaccine Immunity? Why Immunological Memory to Pertussis Is Failing. Cold Spring Harb. Perspect. Biol. 9, a029553 (2017).

9 Hall, J. M. et al. Evaluation of immune responses to Tdap booster during pregnancy in women who received whole cell or acellular pertussis vaccines during childhood. Emerg. Microbes Infect. 14, 2547732 (2025).

10 da Silva Antunes, R., et al. Th1/Th17 polarization persists following whole-cell pertussis vaccination despite repeated acellular boosters. J. Clin. Invest. 128, 3853–3865 (2018).

11 Warfel, J. M., Zimmerman, L. I. & Merkel, T. J. Acellular pertussis vaccines protect against disease but fail to prevent infection and transmission in a nonhuman primate model. Proc. Natl Acad. Sci. USA 111, 787–792 (2014).

12 Creech, C. B. et al. Immunologic profiling of the infant immune response to whole-cell and acellular pertussis vaccines. npj Vaccines 10, 189 (2025).

13 Burdin, N., Handy, L. K. & Plotkin, S. A. What Is Wrong with Pertussis Vaccine Immunity? The Problem of Waning Effectiveness of Pertussis Vaccines. Cold Spring Harb. Perspect. Biol. 9, a029454 (2017).

14 Skoff, T. H. & Martin, S. W. Impact of Tetanus Toxoid, Reduced Diphtheria Toxoid, and Acellular Pertussis Vaccinations on Reported Pertussis Cases Among Those 11 to 18 Years of Age in an Era of Waning Pertussis Immunity: A Follow-up Analysis. JAMA Pediatr. 170, 453–458 (2016).

15 Lefrancq, N. et al. Global spatial dynamics and vaccine-induced fitness changes of *Bordetella pertussis*. Sci. Transl. Med. 14, eabn3253 (2022).

16 Li, Z. et al. Genomic epidemiology and evolution of *Bordetella pertussis* under the vaccination pressure of acellular vaccines in Beijing, China, 2020-2023. Emerg. Microbes Infect. 14, 2447611 (2025).

17 Dai, H. et al. Pathogenicity and vaccine protection of circulating pertactin- and filamentous hemagglutinin-deficient *Bordetella pertussis* strains. Emerg. Microbes Infect. 15, 2640283 (2026).

18 Feng, Y. et al. Emerging macrolide resistance in *Bordetella pertussis* in mainland China: Findings and warning from the global pertussis initiative. Lancet Reg. Health West Pac. 8, 100098 (2021).

19 Xu, Z. et al. Genomic surveillance reveals global spread of macrolide-resistant *Bordetella pertussis* linked to vaccine changes. J. Clin. Microbiol. 63, e0106425 (2025).

20 Huang, L. et al. Rational optimization of glycoprotein E (gE)-encoding mRNA for improved Varicella-zoster virus mRNA vaccine development. Emerg. Microbes Infect. 13, 2392661 (2024).

21 Huang, L. et al. Herpes zoster mRNA vaccine induces superior vaccine immunity over licensed vaccine in mice and rhesus macaques. Emerg. Microbes Infect. 13, 2309985 (2024).

22 Zhao, H. et al. A therapeutic hepatitis B mRNA vaccine with strong immunogenicity and persistent virological suppression. npj Vaccines 9, 22 (2024).

23 Hellgren, F. et al. Unmodified rabies mRNA vaccine elicits high cross-neutralizing antibody titers and diverse B cell memory responses. Nat. Commun. 14, 3713 (2023).

24 Liang, J. L. et al. Prevention of Pertussis, Tetanus, and Diphtheria with Vaccines in the United States: Recommendations of the Advisory Committee on Immunization Practices (ACIP). MMWR Recomm. Rep. 67, 1–44 (2018).

25 Kamachi, K. & Arakawa, Y. Expression of a C terminally truncated form of pertussis toxin S1 subunit effectively induces protection against pertussis toxin following DNA-based immunization. Infect. Immun. 72, 4293–4296 (2004).

26 Di Tommaso, A. et al. Identification of subregions of Bordetella pertussis filamentous hemagglutinin that stimulate human T-cell responses. Infect. Immun. 59, 3313–3315 (1991).

27 Leininger, E. et al. Immunodominant domains present on the *Bordetella pertussis* vaccine component filamentous hemagglutinin. J. Infect. Dis. 175, 1423–1431 (1997).

28 Asgarian-Omran, H. et al. Restricted antibody response to *Bordetella pertussis* filamentous hemagglutinin induced by whole-cell and acellular pertussis vaccines. Infect. Dis. (Lond.) 48, 127–132 (2016).

29 Zhang, H. et al. Algorithm for optimized mRNA design improves stability and immunogenicity. Nature 621, 396–403 (2023).

30 Shao, X. et al. Leveraging an mRNA Platform for the Development of Vaccines Against Egg Allergy. Vaccines (Basel) 13 448 (2025).

31 Guo, K. et al. A novel ionizable lipid with comprehensive improvements in transfection potency, immune profile and safety of lipid nanoparticle. J. Control. Release 386, 114126 (2025).

32 Wang, G. et al. A dual mRNA therapy for treating rheumatoid arthritis and osteoarthritis by inhibiting inflammation and preventing cartilage degeneration. Mol. Ther. (2026).

33 Taranger, J. et al. Correlation between pertussis toxin IgG antibodies in postvaccination sera and subsequent protection against pertussis. J. Infect. Dis. 181, 1010–1013 (2000).

34 Esposito, S. et al. Pertussis Prevention: Reasons for Resurgence, and Differences in the Current Acellular Pertussis Vaccines. Front. Immunol. 10, 1344 (2019).

35 Wang, R., et al. The interaction of innate immune and adaptive immune system. MedComm 5, e714 (2024).

36 Ross, P. J. et al. Relative contribution of Th1 and Th17 cells in adaptive immunity to *Bordetella pertussis*: towards the rational design of an improved acellular pertussis vaccine. PLoS Pathog. 9, e1003264 (2013).

37 Abu-Raya, B. et al. Vaccination in Pregnancy against Pertussis: A Consensus Statement on Behalf of the Global Pertussis Initiative. Vaccines 10 1990 (2022).

38 Warfel, J. M., Papin, J. F., Wolf, R. F., Zimmerman, L. I. & Merkel, T. J. Maternal and neonatal vaccination protects newborn baboons from pertussis infection. J. Infect. Dis. 210, 604– 610 (2014).

39 Hu, Y. et al. Sharp rise in high-virulence *Bordetella pertussis* with macrolides resistance in Northern China. Emerg. Microbes Infect. 14, 2475841 (2025).

40 Mir-Cros, A. et al. Pertactin-Deficient *Bordetella pertussis* with Unusual Mechanism of Pertactin Disruption, Spain, 1986-2018. Emerg. Infect. Dis. 28, 967–976 (2022).

41 Liu, J., Lu, G. & Qiao, J. Global resurgence of pertussis in infants. BMJ 391, r2169 (2025).

42 Amor, N. P. et al. Lipid Nanoparticle: Beyond Delivery Vehicle-Unveiling Its Immunological Adjuvant Potential. FASEB J. 39, e70641 (2025).

43 Verbeke, R., Hogan, M. J., Lore, K. & Pardi, N. Innate immune mechanisms of mRNA vaccines. Immunity 55, 1993–2005 (2022).

44 Kapil, P. & Merkel, T. J. Pertussis vaccines and protective immunity. Curr. Opin. Immunol. 59, 72–78 (2019).

45 Rowe, J. et al. Th2-associated local reactions to the acellular diphtheria-tetanus-pertussis vaccine in 4- to 6-year-old children. Infect. Immun. 73, 8130–8135 (2005).

46 Baptista, P. N., Magalhaes, V. S. & Rodrigues, L. C. The role of adults in household outbreaks of pertussis. Int. J. Infect. Dis. 14, e111–e114 (2010).

47 Ivaska, L., Barkoff, A. M., Mertsola, J. & He, Q. Macrolide Resistance in *Bordetella pertussis*: Current Situation and Future Challenges. Antibiotics 11 1570 (2022).

48 Taniguchi, K. et al. Clinical and Microbiological Characteristics of macrolide-resistant *Bordetella pertussis* Infection: A case series in Osaka, Japan (2024-2025). J. Infect. Chemother. 32, 102916 (2026).

49 Tsukahara, H. et al. Severe macrolide-resistant *Bordetella pertussis* (ptxP3) infection in Japanese infants: First report of cases requiring intensive care. Int. J. Infect. Dis. 158, 107960 (2025).

50 Wang, D. et al. mRNA-LNP delivery of a homo-tandem nanobody protects goslings from novel goose astrovirus infection. J. Virol. 100, e0112726 (2026).

