## Supplemental information for "A tricomponent mRNA pertussis vaccine elicits superior protective immunity over licensed acellular pertussis vaccine against emerging antibiotic-resistant variants"

### Extended Data Figure 1

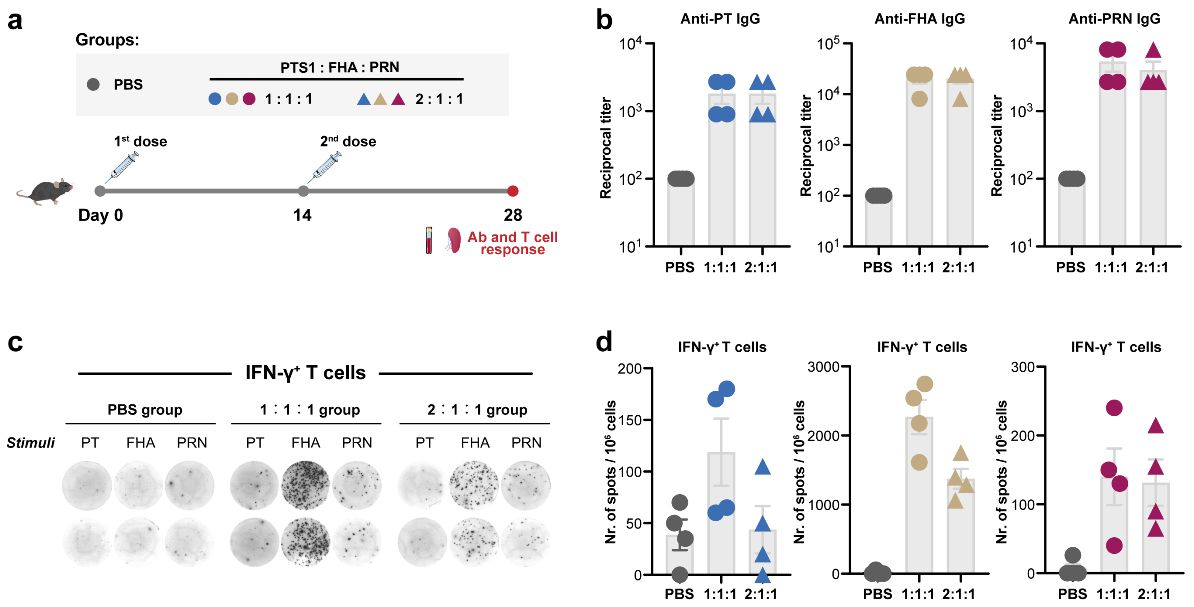


**Extended Data Figure 1. Comparison of Immunogenicity of two different tricomponent mRNA pertussis vaccine formulations.**

**a,** Experimental design schematic. C57BL/6 mice (n=4/group) were i.m administered with two different mRNA pertussis vaccine formulations at a dose of 10 μg following a prime-boost regimen. **b,** Antigen-specific IgG at day 14 after boost immunization was evaluated. Endpoint titers are shown. **c,** Splenocytes were stimulated with inactivated PT, FHA, or PRN (each at 2 μg/mL) for 20 hours. Frequencies of IFN-γ producing T cells were measured by ELISpot assay. Data from two representative animals are shown. **d,** Numbers of antigen-specific T cells were enumerated and indicated as spots per million stimulated cells. Data are presented as mean ± SEM.

### Extended Data Figure 2

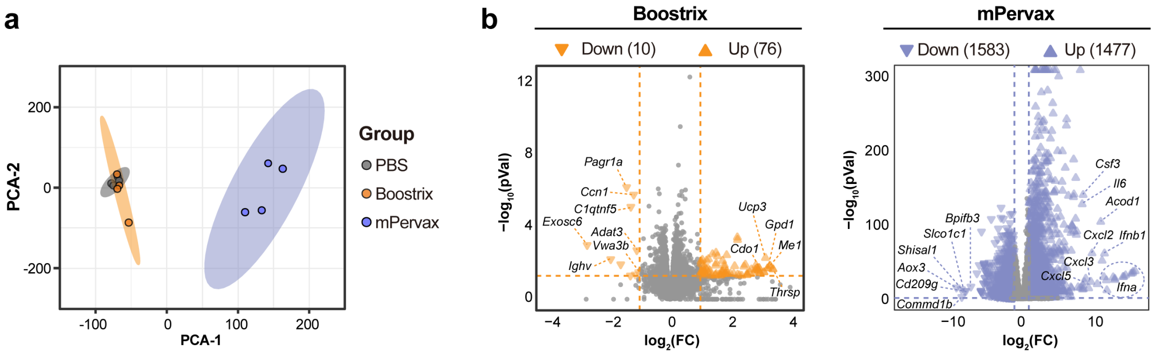


**Extended Data Figure 2. Transcriptomic analysis of mPervax in comparison with Boostrix.** Mice were treated following the experimental schedule described in Figure 2a. 12 hours post injection, dLNs (4 biological replicates/group) were collected and subjected to bulk RNA sequencing. **a,** Principal component analysis (PCA) of global gene expression profiles. **b,** Volcano plots illustrating differentially expressed genes (DEGs) for each treatment group compared to the PBS group. DEGs were identified using the criteria of $p < 0.05$ and |fold change| ≥ 2.

**Extended Data Figure 3**

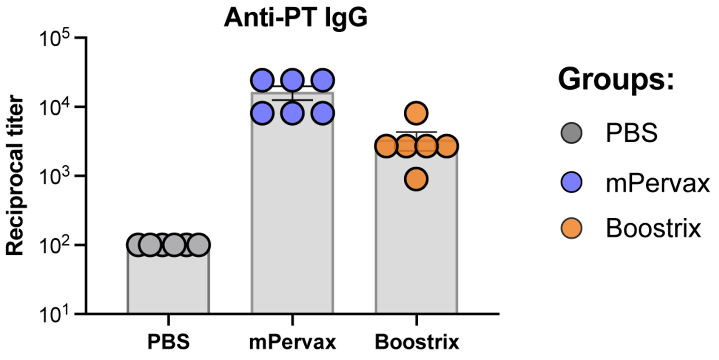


**Extended Data Figure 3. Measurement of anti-PT IgG titers.**

Mice were treated following the experimental schedule described in Figure 5a. At day 28, serum samples were analyzed for anti-PT IgG titers. Endpoint titers are shown. Data are presented as mean ± SEM.

**Extended Data Figure 4**

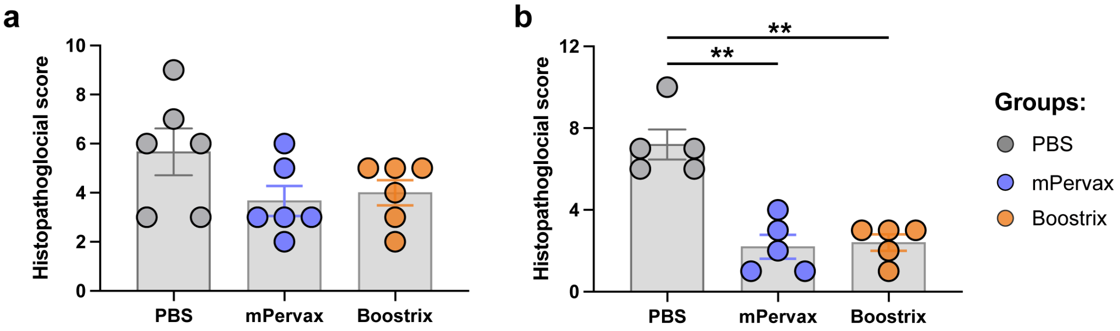


**Extended Data Figure 4. Histopathological scores of the lung tissues.**

Mice were treated following the experimental schedule described in Figure 5a. Seven days post bacterial challenge, animals were necropsied and lung specimens were subjected to H&E staining for histopathological examination. **a,** Histopathological scores of lung tissues corresponding to Figure 5c. **b,** Histopathological scores of lung tissues corresponding to Figure 5f. Data are presented as mean ± SEM. One-way analysis of variance (ANOVA) with Tukey's post hoc test was used for statistical comparison. $^{**}p < 0.01$.

**Extended Data Figure 5**

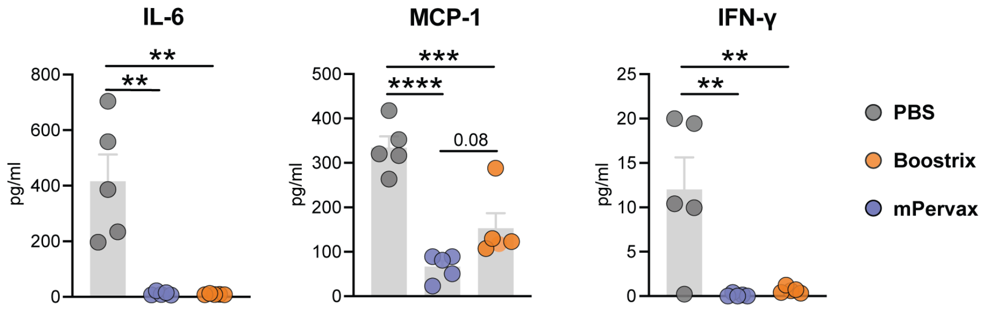


**Extended Data Figure 5. Measurement of serum inflammatory cytokines.**

Mice were treated following the experimental schedule described in Figure 5d. Seven days post bacterial challenge in the offspring mice, serum samples were collected for evaluation of IL-6, MCP-1 and IFN-γ using BD® Cytometric Bead Array (CBA). Data are presented as mean ± SEM. One-way analysis of variance (ANOVA) with Tukey's post hoc test was used for statistical comparison. ***p* < 0.01, ****p* < 0.001, *****p* < 0.0001.
